# A Randomized Trial of Grant Writing Coaching Groups: Qualitative Interviews Revealing Key Elements of Intervention Efficacy

**DOI:** 10.64898/2026.09.17.752282

**Authors:** Julia Mendes, Christine Wood, Remi Jones, Jeffrey Engler, Anne Marie Weber-Main, Harlan Jones, Melanie Steiner, Kola Okuyemi, Richard McGee

## Abstract

**Background:** Training in grant proposal writing is an essential component of professional development for academic scientists in the biomedical and behavioral sciences. Despite the expansion of inter- and intra-institutional grant writing coaching groups as an approach to honing these skills, specific features that enhance or limit coaching group effectiveness have not been rigorously studied.

**Methods:** Qualitative interviews were conducted with a subset of early-career investigators (n=204 and coaches (n=36) engaged in a national, U.S. based, group-randomized trial of grant writing coaching groups to test the effects of two variables on submission and funding of national-level proposals: (1) coaching duration (regular/extended dose) and (2) mode of engaging a scientific advisor (someone with content-aligned expertise) in the coaching process. This report focuses on interviews conducted upon completion of the regular coaching dose – 5 months of biweekly, group-based coaching sessions to support active proposal writing. Interviews were designed to identify which coaching group elements were perceived to be the most critical. Transcribed interviews were analyzed using deductive (participants) or open (coaches) coding to identify themes.

**Results:** Triangulation of results from participant and coach interviews showed strong concordance that coaches, peers, and scientific advisors all played key roles in supporting intervention efficacy. Sufficient alignment of scientific fields and/or methodologies among group members was important, although breadth of perspectives was also valued. Other critical group features were skilled and well-organized coaches, detailed feedback, peer-to-peer support (technical and psychosocial), and clear expectations for group functioning. Factors attenuating impact included variation in participants’ engagement and “readiness to write,” within-group mismatches of expertise or grant mechanisms, and limited research support at some participants’ home institutions.

**Conclusion:** This study identified key elements of successful grant writing coaching groups and potential barriers to their effectiveness, while yielding insights about tailoring this approach for individuals at different stages of proposal development.

## Introduction

Securing extramural research funding is a pivotal step in the professional development of early-stage investigators in the biomedical and behavioral sciences. Historically, trainees have acquired grant writing skills through mentorship during their doctoral and postdoctoral training, combined with self-learning and through trial and error. As research training gradually became more structured, grant writing workshops were created as an approach to complementing the experiential knowledge that mentors are expected to provide. Workshop facilitators can offer useful concepts and information about grant writing and review, show examples of successful proposals, and share insights from experienced reviewers. However, short-term training models such as one-time workshops are unable to provide sustained, contextualized engagement with successful writers, and the individualized critical feedback on a participant’s in-progress proposal that are essential for skill maturation in this complex writing genre.

The biomedical research community began to recognize the need for more intensive processes to teach grant writing skills, but such interventions were not available in most training environments until recently. Today, an internet search for grant writing mentoring and coaching reveals that many institutions, particularly high-resourced universities, are providing structured, more intensive grant writing support for early-career scientists. The design of these approaches varies: some provide didactic lectures along with feedback on proposals [1,2] while others focus primarily on feedback while writing [3], and the duration ranges from half a day to six months or longer. In response to this growth of and variations in approaches, investigators in the National Research Mentoring Network (NRMN), funded by the National Institutes of Health (NIH) from 2014 to 2025 [4], developed and evaluated several models of grant writing coaching groups.

In the first phase of NRMN (2014-2019), six variations of national grant writing coaching programs (GCPs) were implemented and evaluated [5,6]. Their designs varied in accordance with their intended audience. For example, one GCP targeted individuals with substantial research experience and publications but who had not yet achieved success with NIH career development (K) or research (R) awards, particularly R01-level grants; participants in this program were ‘ready to write’ and likely able to submit a proposal by the end of a three-month coaching period. Other GCPs were designed for individuals just beginning to formulate plans for a proposal and/or were at lower-resourced institutions with limited support for research development. Despite their differences, all models were similar in that they engaged cohorts of early-career faculty or postdoctoral fellows who met in small groups, often weekly or biweekly, during active proposal development, under the guidance of grant writing coaches (typically faculty with high levels of skills and success with grant writing and review). An evaluation of proposals submitted and funded across the six NRMN GCP variations affirmed the value of the core intervention (i.e., group-based coaching during active phases of proposal writing). However, questions remained as to which intervention features mattered most and the ‘mechanisms’ by which the GCP approaches were beneficial [6].

The second phase of NRMN (2019-2025) funded independent research projects to test hypotheses related to mentoring and professional development, including ideas that emerged from the programmatic efforts of NRMN phase 1. We drew on the original GCP data to create an optimized five-month long, eight-session group coaching intervention for participants who were identified as being sufficiently ‘ready to write’ such that submission of a proposal was a realistic possibility within a year of coaching initiation. However, two key variables remained unclear: (1) the optimal duration of coaching for maximum effectiveness, and (2) whether the involvement of a scientific advisor – someone with expertise in the participant’s specific research area – would be more impactful when integrated into the coaching process than when providing independent advising.

With funding from NRMN phase 2, we tested these two variables in a 2×2 factorial, group-randomized trial that compared four variations of the coaching intervention. Protocol details and an analysis of baseline data from the trial have been published [7,8]. A separate publication (under review) reports quantitative outcomes from the study – specifically, the rates of national-level proposal submissions and awards across study arms and by participants’ demographic and professional characteristics [9]. An overarching finding from those analyses is that there was a high rate of proposal submissions across all four arms. Some arm-specific differences in grant awards were observed, although the effects were nuanced (with subtleties by demographic group and grant mechanism). A clear impact of the writing group design was a substantially higher funding success rate among faculty submitting R01 and R01-like proposals than rates published by NIH over a comparable period of time [9]. The qualitative data reported in this article provide insights into how and why the coaching group model is effective.

This report utilizes data from extensive qualitative interviews with participants and coaches that were conducted one to two months after the completion of the five months of bi-weekly grant writing coaching groups. The objectives of these qualitative analyses were to elucidate (1) elements of the coaching groups that were perceived to be most valuable and effective; (2) the roles of coaches, scientific advisors, and peers in the group coaching process; and (3) differences among individuals that were likely contributors to the success of submitted proposals.

## Methods

### Study overview

The study was initially based at the University of Utah, then transitioned to Indiana University without disruption in 2024 when the Principal Investigator (author KO) accepted a new position. The study was supported by an NIH U01 cooperative agreement award (grant number U01-GM132366) and received an exemption determination by the Institutional Review Board at the University of Utah (IRB number 00113440). Indiana University affirmed this determination.

The study protocol has been previously published [7]. A summary of key details is provided here for context. Participants were recruited for six distinct cohorts between October 2019 and June 2022 from a US-based population of early-career biomedical and behavioral scientists (junior faculty and postdoctoral fellows) through announcements from NRMN. Participants were selected for inclusion based on (1) their status as early-stage investigators; (2) having never received an R01-type grant; (3) having sufficient research training and preliminary data to support the development and submission of a K- or R-type proposal within 12 months; (4) being employees of an institution with sufficient resources to support the proposed research; and (5) having sufficient time to devote to grant writing over the subsequent five months (6). In response to the NRMN phase 1 finding that individuals writing proposals without content expert support were less likely to succeed at submitting them, this NRMN phase 2 study required that prospective participants, at the time of application, identify a Scientific Advisor (SA) with expertise in their research area to assist with proposal development. In essence, participants and SAs applied as a pair.

Previous work has shown that individuals in grant writing groups need to be working in broadly similar fields to provide meaningful feedback [6]. Thus, coaching groups were created based on four categories: (1) basic science/laboratory research; (2) social science/behavioral research; (3) clinical research, including clinical trials; and (4) epidemiology/data science research. Groups included one coach and four to six participants. Among those who applied for each cohort and met the inclusion criteria, individuals were randomly assigned to a group type that best fit their research until a cohort was filled. A coach with expertise in that type of research was then assigned to the group. These groups became the unit of randomization for our study. The study oversampled from the qualified applicant pool for men, low-resourced institutions, and for individuals from racial and ethnic groups historically underrepresented in the biomedical sciences. This was done to achieve sufficient representation from all demographic groups and to allow for statistical comparisons among them.

Following a virtual kick-off meeting introducing the fundamentals of grant writing, coaching groups met virtually for eight two-hour sessions over five months. Each session focused on one major component of an NIH-style grant (e.g., Specific Aims page), and participants were expected to develop and write that section between virtual meetings and upload it to a cloud-sharing site where peers and the coach reviewed it. The group activity also included a mock study review session where participants selected an external subject matter expert to provide feedback on their completed proposal in the style of an NIH reviewer. The timing and focus for each of the group sessions is provided in Table S1.

The study was designed as a pragmatic, unblinded, 2×2 factorial, group-randomized trial with four experimental arms to test the impact of two intervention features on proposal submission and funding success: (1) the duration of coaching support (extended versus regular dose) and (2) mode of engaging SAs in the group coaching process (structured versus unstructured). All study arms included a five-month period of coaching group engagement. The “regular dose” ended after group coaching; the “extended dose” provided participants with continued access to their coach for an additional 18 months of one-on-one interaction as desired. The conditions of SA engagement tested an “unstructured” model, where SAs worked with participants outside of the coaching group meetings on a mutually determined basis (no contact between the SA and coach), and a “structured” model, where SAs had multiple contact points with the coaching process (i.e., SAs provided the coach with a written review of an early draft of the participant’s Specific Aims page, attended at least one coaching group meeting during the five-month duration, had one separate meeting with the participant and coach together, and optionally joined the mock review). Other meetings and feedback between participants and SAs were at their discretion. A description of the four arms is provided in Table 1, and the timing of the six cohorts starting in January 2020 is shown in Table 2. The study spanned the COVID-19 pandemic and its immediate aftermath.

**Table 1.** Study Arms.

|  |  |
| --- | --- |
| • **Regular Dose** (5-month of group coaching)<br>• **Unstructured** (SA not engaged with coaching group or Coach) | • **Extended Dose** (18 months of access to Coach following Regular Dose)<br>• **Unstructured** |
| • **Regular Dose**<br>• **Structured** (SA joins coaching group once and meets once with Coach + participant) | • **Extended Dose**<br>• **Structured** |

The table describes how regular dose coaching vs. extended dose coaching, and inclusion of scientific advisors with the group coaching process, were distributed in the 4 arms.

**Table 2.** Cohort Schedule 2020-2024.

|  | Jan 2020 | Jun 2020 | Jan 2021 | Jun 2021 | Jan 2022 | Jun 2022 | Jan 2023 | Jun 2023 | Jan 2024 |
| --- | --- | --- | --- | --- | --- | --- | --- | --- | --- |
| Cohort 1 | Regular Dose (5 months) | Extended Dose (18 months) |  |  |  |  |  |  |  |
| Cohort 2 |  | Regular Dose | Extended Dose |  |  |  |  |  |  |
| Cohort 3 |  |  | Regular Dose | Extended Dose |  |  |  |  |  |
| Cohort 4 |  |  |  | Regular Dose | Extended Dose |  |  |  |  |
| Cohort 5 |  |  |  |  | Regular Dose | Extended Dose |  |  |  |
| Cohort 6 |  |  |  |  |  | Regular Dose | Extended Dose |  |  |

The table illustrates the starting date for each cohort with the timing of the group coaching and subsequent access to the coach for the extended dose.

### Coaches

Coaches were recruited among faculty based on their status as established investigators with prior NIH funding and with experience as reviewers on NIH study sections. Most coaches had some prior experience leading grant writing groups through NRMN or other programs. New coaches were also recruited to limit the number of coaches leading groups in more than one cohort. Recommendations for new coaches were made by experienced coaches based on knowledge of their success obtaining grants, experience reviewing NIH-style proposals, and efforts with research training. All coaches were provided with a coaching manual with a description of the general method of group coaching being employed and specifics of their roles in the study (7). Additionally, new coaches were trained virtually in the coaching model and were observed by one of the study leaders during their first coaching session. New coaches quickly and effectively understood and adopted the group coaching model and mirrored the approaches of experienced coaches.

### Recruitment of participants and coaches for interviews

The study sample included 367 participants (271 faculty and 96 postdoctoral fellows) across six cohorts, with four arms in each cohort. All participants were invited to complete an interview soon after the initial five-month coaching groups ended. Of those invited, 204 participants (56%) completed an interview. Analysis of the demographic characteristics of those who accepted the interview invitation showed that they were demographically indistinguishable from the total sample with respect to gender and race and ethnicity.

Of the 57 coaches who coached one or more groups, 36 completed at least one interview soon after finishing one of the five-month coaching groups; three coaches completed multiple interviews because they coached in more than one cohort.

### Interviews with participants and coaches

Interview guides are provided as Table S2. Interviews with participants collected information about the strengths and value of coaches; the structure of the group meetings; challenges with the coaching group; components of the program such as the mock review; and the quality of feedback received from coaches and peers. Participants were also invited to complete an interview at the end of 24 months from the start of their cohorts to learn more about what transpired with or without additional access to coaches (the extended dose condition). Those interviews will be the subject of future publications.

Interviews with coaches collected information about their skillsets and contributions to the groups; the strengths and limitations of the group; the structure of meetings; experiences with program elements such as the mock review; and suggestions for future iterations of coaching groups. Coaches from the extended dose arms were also invited to complete interviews after 24 months. Those interviews will be the subject of future publications.

All interviews were conducted by three experienced qualitative researchers at Northwestern University who were not involved with the study in any other way. Interviewers were familiar with the training of biomedical scientists through their involvement with the National Longitudinal Study of Young Life Scientists [10]

### Interview coding and analysis

The participant interview guide was semi-structured. The coding architecture was constructed deductively based on the topics of the questions and formed into initial primary themes for directed content analysis [11]. The coach interview guide was more open-ended than the participant interview guide. Analysis of the coach interviews included open coding to inductively identify themes followed by constant comparison across interviews to finalize themes [12]. In all cases, interviewers had the ability to ask follow-up questions that probed for more detailed information and broadened the types of responses.

Initial coding of the participant interviews was done by one of the authors (JM), with regular review and concurrence with one of the co-investigators of the study (RM) to provide additional context. After initial coding, sub-codes were created to capture more granular commonalities of themes. All sub-coding was reviewed by JM and RM to arrive at final themes. Another author (CW) completed coding of the coach interviews, with regular review and concurrence with the study co-investigator (RM). Coding was completed using Dedoose qualitative data analysis software.

Qualitative analysis is more time-consuming than quantitative assessment of outcomes, such as the number of proposals submitted and funded. It was not practical to delay qualitative analysis until after the last cohort and still complete it during the period of funding. Therefore, the decision was made to start with a complete qualitative analysis of the first three cohorts. Of the 204 interviews with participants across six cohorts, the data in this report are drawn from 119 individuals from cohorts one through three. These cohorts contained 168 participants. Thus, the data in this paper include a very high fraction (71%) of individuals in these cohorts. To check for reliability and generalizability across cohorts four through six, coding and analysis of a subset of interviews from cohort five, which began in January 2022, were conducted. No systematic or substantive differences between cohort 5 and cohorts 1-3 were observed. Therefore, we consider the first three cohorts to be representative from a methodological and findings perspective to the subsequent cohorts.

Twenty-six of 29 coaches from cohorts 1-3 completed an interview (90%). Analysis of the data from those interviews showed no substantive differences from the 10 coaches interviewed in later cohorts. Because the total number of interviews was much smaller, the data presented are from all 39 coaches interviewed across the six cohorts.

## Results

### Interviews with participants

The number of individuals who made comments related to each theme, from the 119 coded interviews, is indicated in parentheses. A summary of the themes from participants is provided in Table 1. Quotations exemplifying these themes are provided in Table S3.

**Table 1.** Summary of Themes from Participant Interviews. The numbers in parentheses indicate the number of individuals who mentioned something fitting within that theme.

| Coaches |  |
| --- | --- |
| Attributes of Effective Coaching | Attributes of Ineffective Coaching |
| Experience and Expertise (45)<br>Coaches' extensive experience in grant writing and reviewing was widely valued<br>Participants appreciated their coaches' deep understanding of what makes a successful proposal<br>Coaches offered nuanced advice on both technical and strategic aspects of grant writing<br>Their expertise boosted participants' skills and confidence in submitting competitive grants | Disciplinary Mismatch (16 mentions):<br>Coaches lacked expertise in participants' specific fields.<br>Feedback was often too general and lacked scientific depth.<br>Misalignment reduced the relevance and impact of coaching. |
| Real World Relevant Feedback (34)<br>Feedback reflected real-world grant review standards and expectations<br>Coaches simulated the scrutiny of actual grant reviewers, better preparing participants for submission<br>Specific, detailed input on grant mechanisms and strategic alignment was especially helpful<br>Coach feedback mirrored what might be heard in a study section | Vague or Unhelpful Feedback (10)<br>Feedback lacked specificity and actionable suggestions<br>General comments on clarity or structure were not detailed enough<br>Participants were left uncertain about how to revise their proposals<br>This was often linked to disciplinary unfamiliarity or vague guidance |
| Quality of Advice and Mentoring (32)<br>Coaches provided honest, critical, and constructive feedback in a supportive way<br>Participants valued the assertive yet nurturing mentoring style<br>Honest critiques were essential for participant growth and improvement<br>Coaches' objectivity (being outside participants' local institutions) ensured unbiased feedback<br>External perspective helped ground advice in broader grantsmanship principles<br>Coaches shared helpful materials like successful grant examples and tools | External Stressors Affected Coaches (9)<br>Coaches were impacted by external pressures, especially during COVID-19.<br>Personal and professional stress limited their availability and focus<br>Some coaches appeared distracted or overwhelmed during sessions<br>These issues reduced the overall effectiveness of the coaching |
| Organization and Preparation (23)<br>Coaches structured the process with clear, detailed meeting plans and timelines<br>Tasks were well-defined<br>Organized meeting flow and consistent scheduling supported steady, focused progress<br>Participants credited this structure with helping them systematically address their proposals | Ineffective Communication (7)<br>Feedback was unclear or lacked structure<br>Coaches seemed unprepared for meetings<br>Poor communication left participants unsure of next steps<br>Sessions felt less productive |
| Supervision (19)<br>Coaches actively supervised the peer review process and mediated disputes<br>They clarified confusion when peer feedback was unclear or contradictory<br>Having a coach as the final authority added reliability and built trust<br>Participants felt confident deferring to coaches when navigating mixed opinions | Lack of Specific Feedback on Grantsmanship (6)<br>Strategic and technical aspects of grantwriting were overlooked<br>Feedback focused on general writing rather than proposal development<br>Participants wanted more training on grant mechanics and strategy |
| Approachability and Understanding (16) | Unequal Attention (5) |
| Coaches were described as patient, non-judgmental, and approachable<br>Participants felt comfortable asking questions without fear of judgment<br>A safe, supportive environment fostered open dialogue and better learning<br>This demeanor strengthened mentoring relationships and promoted participant development | Coaches did not distribute attention equally among group members<br>Some participants received minimal guidance, affecting their progress<br>Perceived favoritism harmed group cohesion and morale |
| Cross-Disciplinary Perspectives (14)<br>Feedback from coaches in other disciplines was beneficial<br>Cross-disciplinary insights improved the logic, clarity, and overall presentation of proposals<br>Participants appreciated the ability of coaches to provide relevant feedback, regardless of field<br>This highlighted the coaches' versatility and depth of grant-writing expertise<br>When coaches *did* share the participant's disciplinary background, feedback was seen as especially tailored and insightful<br>Exposure to both aligned and cross-disciplinary perspectives mirrored real grant review panels<br>This broadened participants' thinking and helped make their proposals more accessible to diverse reviewers |  |

| Interactions with Peers in the Groups |  |
| --- | --- |
| Benefits of Peer Interaction | Limitations of Peer Interaction/Feedback |
| Feedback (25)<br>Feedback from different disciplines helped participants view their work from new angles<br>Cross-disciplinary insights revealed overlooked issues and improved proposal clarity<br>Feedback mirrored real grant panel diversity, helping refine communication for broader audiences<br>A clear, organized feedback process improved participants' critique and revision skills<br>Exposure to both high-level and detailed feedback strengthened different proposal sections<br>Peer input helped identify weaknesses and guided meaningful revisions | Low Participation and Turnout (17)<br>Some groups had inconsistent attendance, with only one or two peers showing up regularly<br>Sporadic participation disrupted continuity and weakened group dynamics<br>Peers often failed to share drafts on time, limiting meaningful feedback<br>Some sessions were canceled or lost value due to low engagement |
| Engaging with Peers from Various Career Levels and Backgrounds (24)<br>Participants benefited from interacting with peers at different career stages<br>Early-career researchers gained mentorship from more experienced colleagues | Time Constraints (12)<br>Limited time restricted deep engagement with peer drafts<br>Feedback felt rushed due to the volume of documents and short review windows |
| Varied disciplinary and professional backgrounds brought diverse insights to each proposal<br>The exchange enriched group discussions and deepened understanding of grant-writing | Participants struggled to balance reviewing others' work with progressing on their own<br>Last-minute draft submissions reduced the quality of peer feedback |
| **Support and Camaraderie (23)**<br>Groups fostered emotional support and a non-competitive atmosphere.<br>Peer encouragement helped participants stay motivated during difficult periods, like COVID-19.<br>A sense of community encouraged honest, constructive feedback and collaborative growth. | **Unfamiliarity with Grant Mechanisms (9)**<br>Peers often lacked familiarity with different grant types<br>Peers' ability to provide relevant, mechanism specific feedback was limited<br>Feedback was less useful when participants worked on different types of grants |
| **Sharing of Resources (10)**<br>Participants exchanged tools, templates, and grant-writing materials<br>Shared links to NIH guidelines and sample documents improved everyone's work<br>Resource-sharing expanded participants' grant-writing toolkits and strategies | **Disciplinary Misalignment (9)**<br>Peers unfamiliar with a participant's field gave superficial or grammar-focused feedback<br>Lack of shared methods of research context limited substantive critique<br>Interdisciplinary input was less effective when deeper field-specific understanding was needed |

| **Scientific Advisors** |  |
| --- | --- |
| **Benefits of Scientific Advisor Participation** | **Limitations of Scientific Advisor Participation** |
| **Additional, Insightful, and Helpful Feedback (26)**<br>Advisors gave direct, targeted feedback that improved drafts<br>Helped identify weaknesses, suggest alternatives, and strengthen arguments<br>Shared knowledge of funding landscapes and past grant experiences<br>Provided examples of successful applications as helpful references | **Scientific Advisor Didn't Participate at All (11)**<br>Advisors in the unstructured arm did not engage in group activities<br>Participants lacked guidance on when or how to connect with their advisors<br>Some advisors were invited but didn't attend sessions or respond<br>Participants hesitated to burden busy advisors, leading to minimal interaction |
| **Existing Relationship with Scientific Advisor Improved (16)**<br>Study group involvement enhanced advisor-participant communication<br>Regular meetings fostered accountability and consistent engagement<br>Advisors became more familiar with participants' projects and goals<br>Strengthened mentorship through clearer understanding of research interests | **Scientific Advisors had Busy Schedules and Other Time Commitments (8)**<br>Advisors had limited ability due to other professional responsibilities<br>Scheduling meetings was difficult, and they were often rushed<br>Advisor commitments conflicted with program timeline<br>Participants felt unsupported at key points in the writing process |
| **Scientific Advisor Complemented and Filled in Gaps with Group Coaches (13)** | **Advisors were not Helpful Overall (4)**<br>Some advisors gave minimal or overly general feedback |
| • Advisors and coaches offered distinct, complementary support<br>• Coaches focused on writing strategy; advisors on technical/scientific accuracy<br>• Dual perspectives improved both content quality and communication<br>• Aligned feedback from both roles reassured participants and clarified revisions | • Feedback focused on grammar rather than scientific content or structure<br>• A lack of topic-specific expertise limited their usefulness |
| Expertise and Knowledge on Topics and Grants (12)<br>• Advisors offered field-specific knowledge crucial for niche or complex topics<br>• Guided participants on study sections, literature, and experimental design<br>• Helped align proposals with reviewer expectations and funding priorities<br>• Directed participants to relevant resources and strengthened proposal framing |  |

| Meeting Format and Structure |  |
| --- | --- |
| Strengths | Limitations |
| Accountability and Deadlines (14)<br>• Meeting structure encouraged consistent progress<br>• Deadlines promoted steady writing habits and reduced procrastination<br>• Time constraints helped participants who work better under pressure<br>• Scheduled presentations motivated participants to stay on track despite other academic responsibilities | Different Grant Mechanisms (17)<br>• Participants worked on various grant types, leading to mismatched content and focus<br>• Meetings sometimes felt unproductive when grant-specific needs weren't addressed<br>• Those writing career development grants often felt unsupported due to lack of group expertise in that area |
| Collaborative Learning (13)<br>• Meetings enabled feedback and interaction with peers from diverse disciplines<br>• Peer input offered new perspectives and highlighted areas for improvement<br>• Reviewing others' work enhanced participants' ability to critique and strengthen their own writing | Online Meeting Issues (13)<br>• Time zone differences made scheduling difficult<br>• Technical problems disrupted participation<br>• Lack of in-person interaction reduced engagement and connection |
| Organized and Focused Progression (11)<br>• Meetings focused on specific grant sections in a structured sequence<br>• Systematic approach allowed deep focus on one section at a time<br>• Clear, phased structure guided participants through the grant-writing process logically and manageably | Lack of/Unclear Communication (10)<br>• Inconsistent or unclear communication from organizers<br>• Confusion about deadlines, meeting expectations, and participant roles<br>• Miscommunication occasionally disrupted meeting flow and reduced productivity |
| Supportive Environment (10)<br>• Participants felt safe sharing incomplete drafts and receiving feedback | Too Many Sessions or Meeting too Frequently (8)<br>• Frequent meetings (every two weeks) were overwhelming for |
| • Group meetings built camaraderie and reduced the isolation of grant writing<br>• Emotional and practical support from peers made the writing process feel less intimidating and more collaborative | participants with other professional obligations.<br>• Short turnaround times made it difficult to revise drafts or process feedback.<br>• Participants suggested a three-to-four-week interval to allow for deeper engagement and improved work quality. |

| Mock Review |  |
| --- | --- |
| Mock Review was Helpful | Mock Review was not Helpful |
| Received Valuable Feedback (17)<br>• Mock reviewers provided detailed, constructive criticism<br>• Feedback covered aspects often missed by advisors or coaches<br>• Insights focused on feasibility, significance, and innovation<br>• Helped participants refine arguments and highlight fundable elements | Reviewers were Inadequate (3)<br>• Some reviewers lacked the expertise to give valuable feedback<br>• Reviewers were unresponsive or canceled participation<br>• Feedback was minimal or absent<br>• Participants felt unsupported |
| Good Reviewers and Disciplinary Areas (16)<br>• Selecting experienced reviewers with relevant expertise was key<br>• Reviewers with disciplinary alignment gave more nuanced, targeted feedback<br>• Feedback helped anticipate concerns from actual review panels<br>• Provided guidance on aligning proposals with funding priorities | Feedback Lacked Depth (2)<br>• Feedback was too general or superficial to guide improvements<br>• Critiques focused on broad observations rather than specific proposal issues<br>• Late feedback limited participants' ability to make revisions before submission |
| Improved Grant Writing (11)<br>• Mock review improved overall quality of grant proposals<br>• Highlighted weak areas and gaps in preliminary data<br>• Offered advice on restructuring and clarifying sections<br>• Simulated real study section dynamics, preparing participants for actual review |  |

| Institutional Resources |  |
| --- | --- |
| Sufficient Resources at Home Institution | Insufficient Resources at Home Institution |
| Administrative and Logistical Support (14)<br>• Efficient, well-staffed grants offices that managed deadlines and submissions<br>• Pre-award support teams that helped with budget development and compliance checks<br>• Administrative professionals knowledgeable in NIH and federal grant mechanisms | Insufficient Grant Writing Support Structures (11)<br>• Formal grant-writing support was minimal or absent at some institutions<br>• Workshops were short term and lacked sustained engagement |
| Relief from bureaucratic burdens, allowing researchers to focus on writing | Internal writing groups often included faculty still early in idea development<br>Limited opportunity for meaningful feedback from peers or colleagues |
| Grant Writing Programs and Infrastructure (13)<br>Internal peer review systems that mimicked NIH-style study sections<br>Writing bootcamps and retreats that provided structure and accountability.<br>Access to consultants or specialists who offered strategic feedback.<br>Templates, timelines, and structured feedback cycles to guide the writing process. | Lack of Expertise and Mentorship (7)<br>Difficulty finding mentors with relevant research or grant-writing experience<br>Lack of senior faculty left early-career researchers without role models<br>Participants sought guidance outside their institutions due to internal gaps<br>Limited support for developing funding strategies or long-term research plans |
| Protected Time for Research (12)<br>Teaching release or course reductions to allow for concentrated research time<br>Research-focused faculty appointments with reduced service obligations<br>Institutional policies that prioritized and safeguarded writing time<br>Recognition of grant writing as a core faculty responsibility. | Resource Constraints (7)<br>Inadequate access to research funding, labs, and infrastructure<br>Lack of databases, patient populations, or core facilities slowed progress<br>Insufficient administrative support led to submission errors and delays<br>Participants often had to collaborate with better-resourced institutions |
| Institutional Culture (11)<br>Leadership that visibly valued grant productivity and research excellence<br>Incentives tied to research success (e.g., tenure, promotion criteria)<br>A culture of high expectations balanced with tangible support.<br>Peer communities that modeled and reinforced research-focused behavior | Effect of Institutional Size and Type (4)<br>Smaller or R2 institutions lacked infrastructure and funding capacity<br>Limited internal experience with major grants created a support gap<br>R1 institutions had more resources but often less mentorship support<br>Institutional context shaped both access to resources and culture of collaboration |
| Mentorship and Peer Collaboration (10)<br>Access to experienced, funded faculty for one-on-one mentoring<br>Participation in writing groups or mentoring circles.<br>Peer-to-peer feedback that sharpened writing and built confidence<br>Ongoing support networks that extended beyond formal programs |  |
| Research Facilities and Technical Resources (9) |  |
| • On-campus access to lab space, core facilities, or specialized equipment<br>• Institutional datasets or partnerships that enhanced research feasibility<br>• Technical staff and research coordinators supporting project execution<br>• Infrastructure that enabled participants to demonstrate readiness to funders |  |

| Outcomes Attributed to Participation |
| --- |
| Improved Writing through Various Strategies (24)<br>• Writing became clearer, more concise, and aligned with grant requirements<br>• Participants learned to structure arguments with stronger justification for funding<br>• Emphasis on intentional writing and avoiding redundancy<br>• Adopted strategies to make proposals more reviewer-friendly<br>• Improved time management by spreading writing tasks over time for better quality |
| Better Understanding of Grant Mechanisms (17)<br>• Gained critical insights into the full grant writing process, often for the first time<br>• Learned how to approach key sections like specific aims and research strategy<br>• Feedback and iteration helped avoid common grant-writing mistakes<br>• Improved ability to highlight significance and innovation with reviewers in mind<br>• Developed a stronger grasp of what makes an application competitive |

### Interactions with and contributions of coaches

Participant interviews highlighted a range of coaches’ strengths that contributed to the overall effectiveness of the grant writing groups. The most frequently cited strength was the coaches’ extensive experience in grant writing and peer review (45), which participants credited with enhancing both their technical skills and confidence. Many also praised the real-world relevance of the feedback provided (34), noting that it closely resembled the scrutiny of formal NIH study sections. Participants emphasized the high quality of mentoring (32), describing coaches as honest, supportive, and objective, particularly due to their external institutional status. Strong organizational skills (23), such as clear timelines and structured meetings, were credited with helping participants maintain focus and progress systematically. Coaches also played a key supervisory role (19) by guiding discussions and clarifying conflicting peer feedback. Additionally, their approachability and understanding (16) created a safe environment for learning, and their ability to offer valuable perspectives across disciplines (14) helped participants tailor their proposals for broader audiences. Collectively, these strengths positioned coaches as critical facilitators of both the technical and interpersonal dimensions of grant writing success. As one participant recalled,

> “What I liked most about (my coach) is his extensive experience as a reviewer, having also received R01 grants. I think the balance between what NIH was looking for… (and) I think more of that technical aspect of pulling the proposal together. But he also provided us kind of a balance to that … how to think about the project as your research agenda kind of long-term research goals.”

Despite the many strengths identified in coach performance, a smaller number of individuals brought out limitations that reduced coaching effectiveness in some instances. The most frequently cited issue was a disciplinary mismatch between coaches and participants (16), which led to feedback perceived as being overly general, superficial, or lacking scientific depth. Additional concerns included vague or unhelpful feedback (10), often linked to limited subject matter expertise. Also mentioned were external stressors affecting coach availability and focus, particularly during the COVID-19 pandemic (9). Some participants reported instances of ineffective communication (7), such as unclear expectations or disorganized meetings, and expressed frustration over inadequate guidance on grantsmanship strategy (6). Unequal attention across group members (5), when present, undermined group cohesion and perceived fairness. These findings underscore the importance of aligning coach expertise with participant needs, fostering equitable engagement, and ensuring clarity and preparedness to enhance the overall impact of the GCP.

**Fig 1.**
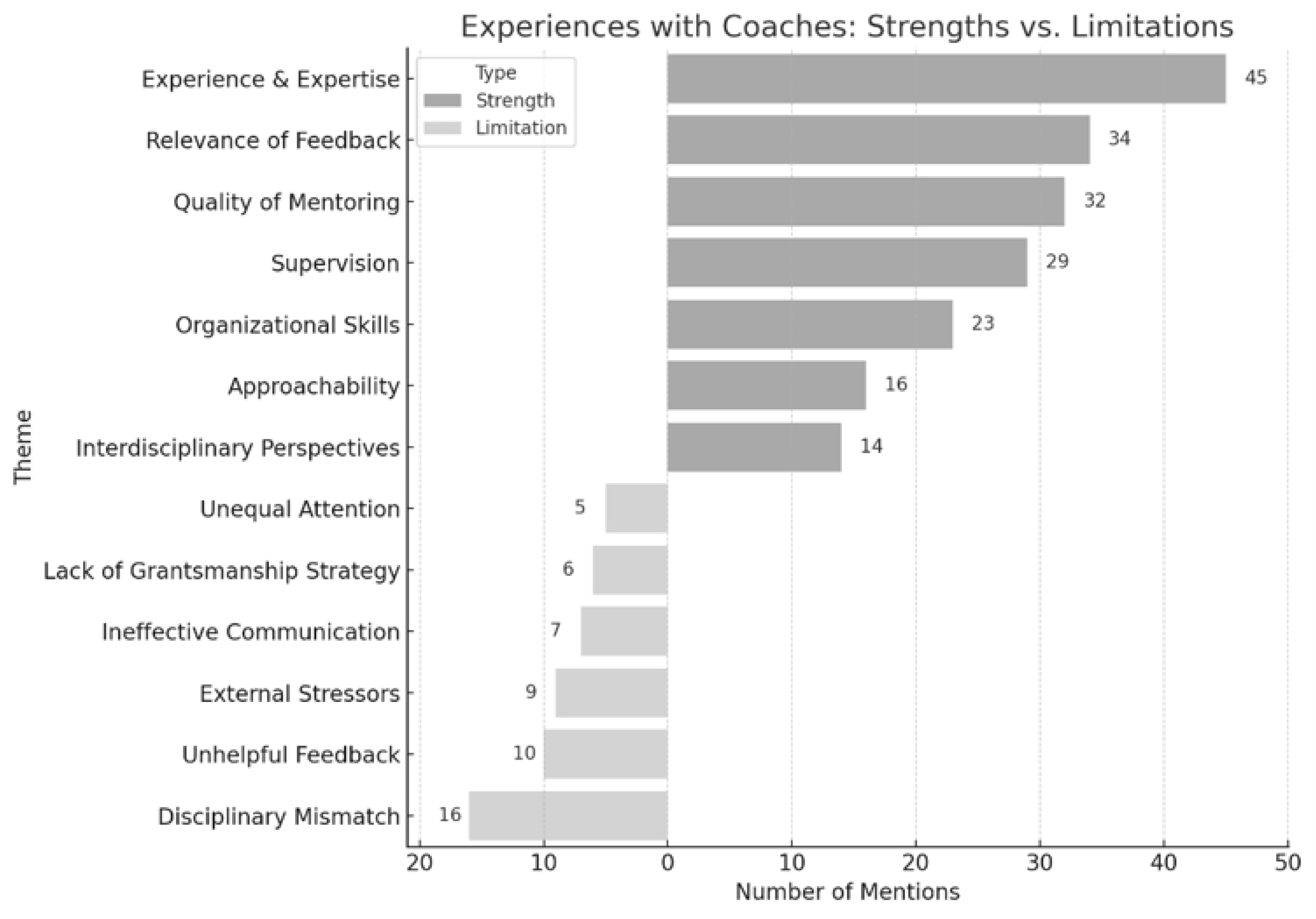
Experiences with Coaches: Strengths vs. Limitations. This figure illustrates participants’ perspectives on the strengths (dark) and limitations (light) of coaches within the grant writing groups.

### Interactions with peers in the groups

Peer interactions were widely viewed as a valuable component of the grant writing groups. Benefits included feedback quality, emotional support, and collaborative learning. Participants emphasized the value of cross-disciplinary feedback (25), engagement with peers across varying career stages (24), and the sense of community fostered through mutual support and encouragement (23). The peer critique processes enhanced revision skills, with resource sharing (10) and ongoing post-group connections (6) contributing to both proposal development and sustained professional networks. For example, one participant noted:

> “It was really helpful to have outside people to keep me accountable and to give me feedback and to support me as I go through that process.”

Despite these strengths, participants reported several barriers that sometimes limited the effectiveness of peer interactions. Challenges included inconsistent attendance and low engagement by some participants (17), time constraints that limited feedback depth (12), and challenges posed by differences in grant mechanisms (9) or disciplinary backgrounds (9). Conflicting peer feedback (9) also complicated the revision process. These findings suggest that while peer support adds significant value, group composition, participation expectations, and time allocation can attenuate the contributions of peers.

**Fig 2:**
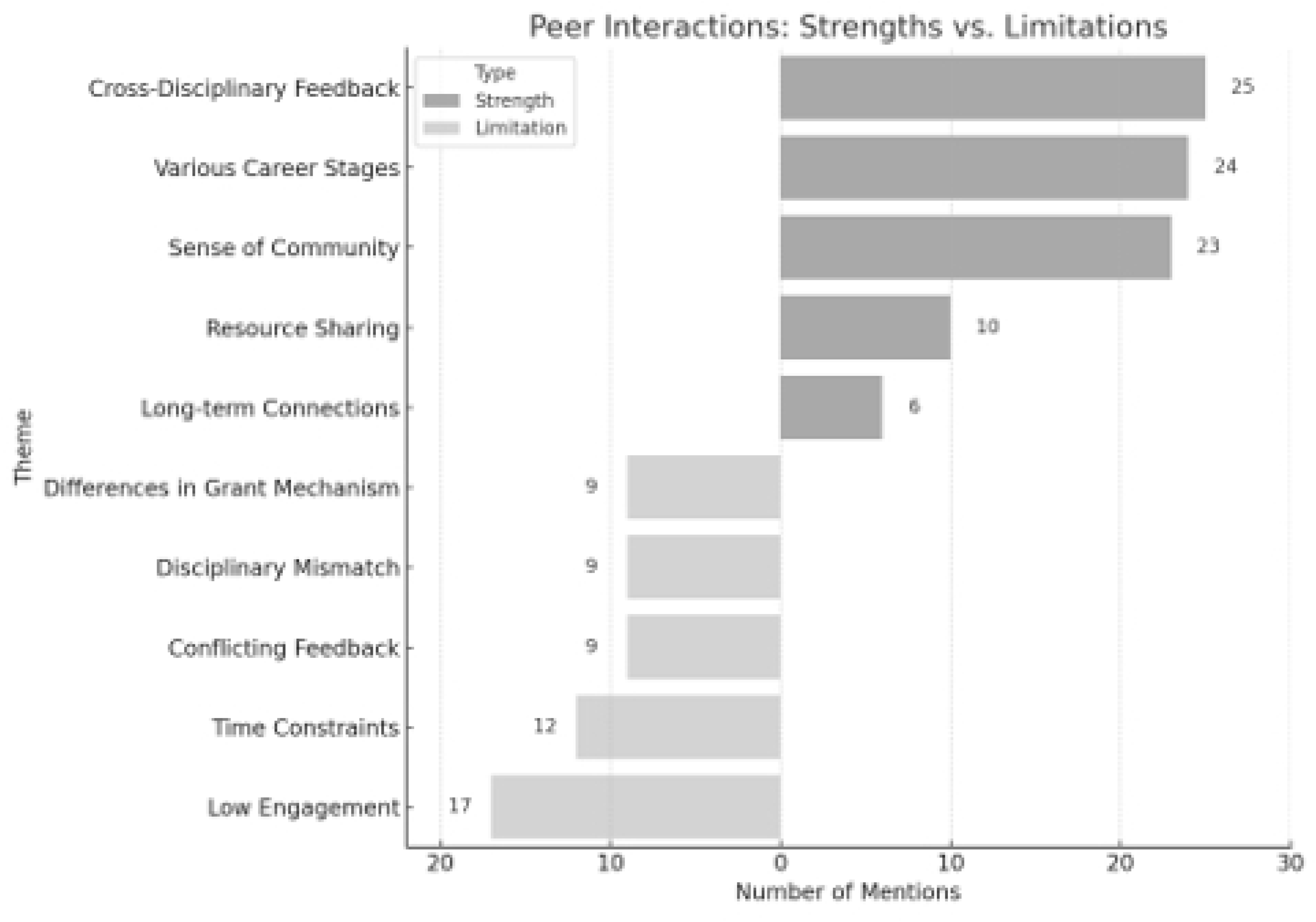
Peer Interactions: Strengths vs. Limitations. This figure summarizes participant feedback on peer interactions, strengths (dark) and limitations (light), within the grant writing groups.

### Interactions with scientific advisors

Most participants selected SAs from their own institutions or existing mentorship networks (14), which facilitated communication but occasionally limited critical feedback. This was particularly true of postdoctoral fellows working on K applications; most of their SAs were either primary or secondary mentors for those proposals. However, some of the participants chose SAs with whom they had no previous relationships. Doing so filled gaps in available scientific expertise in their home institutions and sometimes led to lasting relationships. Overall, participants valued SAs for providing targeted feedback (26), enhancing existing mentorship relationships (16), offering domain-specific expertise (12), and complementing the support of group coaches (13). However, limitations included SA non-participation (11), scheduling conflicts (8), and insufficient or superficial feedback from a few SAs (4).

There were relatively few differences noted by participants on SA involvement between the structured and unstructured arms. For those in the unstructured arm, some reported lack of clarity on how to engage with SAs. This led some participants to take more responsibility for when and how to use the SAs, but sometimes it caused limited engagement. The level and type of engagement between participants and SAs varied more at the individual level than between the structured and unstructured arms.

### Challenges related to the COVID-19 pandemic

The first three cohorts met between January 2020 and mid 2021 during the peak of the COVID-19 pandemic. Analyses of these cohorts showed there were no universal themes attributable to the pandemic, and experiences were different across individual cases. Some participants faced challenges posed by remote work conditions, interruption of clinical trials, and lab closures. These conditions strained some participants and affected their ability to complete the coaching groups. The most frequently mentioned challenges related to the pandemic included generally stressful conditions making it difficult to work (13), delayed and missed deadlines (12), and lab closures delaying acquisition of preliminary data (6). Despite these challenges, few individuals described strong direct impacts on their grant writing and participation in the study. The range of influences was wide, from complete stoppage of progress, indicated above, to freeing up time for writing. Coding and analysis of a subset of interviews from cohort 5, which began in January 2022 after most restrictions had been lifted, showed no systematic or substantive differences from cohorts 1-3, i.e., during or after the primary effects of the pandemic.

### Meeting format & structure

Participants identified both strengths and limitations in the meeting format and structure of the grant writing groups. Commonly praised features included the creation of accountability and clear deadlines (14), opportunities for collaborative learning across disciplines (13), and a structured, sequential approach to writing each grant section (11). Many also highlighted the supportive and collegial environment fostered by regular group meetings (10), which helped reduce the isolation often associated with grant writing. However, several challenges were also reported. The most frequently cited issue was the grouping of participants working on different grant mechanisms (17), which led to content mismatches and unmet needs, particularly for those writing K awards. Other notable concerns included logistical and engagement problems with online meetings (13), unclear communication about expectations (10), overly frequent sessions (8), perceived disorganization (8), and insufficient time for in-depth feedback on grant drafts (7). One participant commented on the frequency of sessions:

> “The meetings were too frequent, and it became overwhelming to keep up. Every two weeks felt like too short a time to make meaningful changes between sessions.”

These findings suggest that while the coaching group format promoted progress and peer support, future iterations may benefit from improved alignment of grant types, clearer communication, and adjusted pacing based on participant workload and group composition.

### Mock review

Only about half of the participants interviewed had a sufficiently complete proposal draft in time for a full mock review. Of those who did, it was widely regarded as a valuable component of the intervention, with 28 participants describing it as helpful. Participants benefited from selecting their own reviewers, which often ensured disciplinary alignment and high-quality feedback. Reviewers provided detailed and constructive critiques (17), particularly on aspects like feasibility, significance, and innovation, which were sometimes missed by coaches or advisors. Reviewers with relevant expertise (16) offered more targeted input, helping participants anticipate reviewer expectations and align proposals with funding priorities. The process also improved overall grant quality (11), highlighted weaknesses, and provided a realistic simulation of NIH-style review. Only a small number of participants reported limitations, generally tied to reviewer selection, timing, or inability to participate due to limited progress.

### Increases in grant writing skills attributed to participation

Many participants reported notable gains in grant writing skills because of their involvement in the study. The most frequently cited improvement was in writing quality and strategy (24), including enhanced clarity, conciseness, alignment with grant requirements, and more effective time management. Many also gained a deeper understanding of grant mechanisms (17), and learned how to structure key sections, avoid common pitfalls, and craft proposals with reviewers in mind. Additionally, some participants noted increased confidence (7), which led to a greater sense of agency, clarity in communicating their strengths, and comfort navigating the grant writing process.

### Institutional resources

Participants’ institutional contexts played a role in shaping their grant writing experiences. Many reported strong institutional support that significantly facilitated their grant writing efforts. Key support included efficient administrative and logistical assistance (14), structured grant writing programs (13), and protected research time through course releases or research-focused appointments (12). Supportive institutional cultures (11), access to experienced mentors and collaborative peer networks (10), and well-equipped research facilities (9) were also noted as critical enablers. These findings highlight the importance of institutional investment in both tangible infrastructure and mentoring systems to support early-career researchers in developing competitive grant proposals.

Some participants described insufficient support at their home institutions, including minimal formal grant writing support (11), limited access to experienced mentors (7), and resource constraints such as inadequate labs, databases, or administrative support (7). These challenges were particularly pronounced at smaller or less research-intensive institutions, where junior faculty may have lacked role models, strategic guidance, or robust collaboration networks. As a result, some participants turned to external mentors or relied on collaborations with better-resourced institutions to bridge these gaps.

### Interviews with coaches

Of 57 coaches, 36 completed at least one interview. Three coaches completed more than one interview and reflected on their experiences across multiple coaching groups. Twenty-seven coaches who completed interviews had prior coaching experience in other programs or interventions. Nine of the new coaches completed interviews.

Coaches had varied views on what was important in performing their roles and the determinants of success for the groups. Predominant themes included: (1) coaching style and skillset; (2) peer support and interaction; (3) SA participation; (4) mock review; and (5) group composition in terms of readiness to write, mix of mechanisms, and mix of disciplines. Analysis revealed that new coaches did not have markedly different perspectives or coaching approaches from experienced coaches; the variation among new coaches was proportional to that among the experienced coaches. Quotations illustrating coach themes appear in Table S4.

### Coach style/skillset

All coaches were asked questions about their skillsets and what they brought to the groups. In many cases, the coaches focused strictly on the development of grant writing skills in their responses. But others described a blended style focused on *both* developing grant writing skills and creating a socially supportive environment for participants. Those focused primarily on grantsmanship discussed their experience of writing grants and acting as reviewers on study sections, and how they used this knowledge to guide participants. Grant writing support included providing detailed feedback on each component of an NIH grant, providing insight into the review process at NIH, and helping to develop skills for writing grants specifically for reviewers on study sections. As one coach reproted:

> “My perspective as a current study section member helped them frame not only their applications in terms of readability but also helped them understand how to sell their ideas to a reviewer or an audience that is not intimately aware of the specific projects or experiments or studies that they’re working on.”

Those with a blended style emphasized the fundamentals of grant writing but found equal value in creating a socially supportive environment where participants could share their experiences without fear of judgment.

### Peer support and interaction

Comments on peer support and interactions appeared in 27 interviews. Coaches commented on peer support in two ways: from a grantsmanship perspective and from a socioemotional perspective. For those focused on grantsmanship, coaches commented on the high-quality feedback group members provided to each other. For example, group members offered support for applying for grants under specific mechanisms and gave detailed feedback on the writing itself. In some cases, participants were able to provide additional subject matter expertise. From a socioemotional perspective, groups provided a sense of camaraderie and community based on shared experiences. “[A] great thing that was valuable to them was the insight that each of the members of the group had for what the others were going through.” There were almost no negative comments about peer interactions, except in cases where groups had low attendance or where members dropped out prematurely. Peer support is a major strength of the coaching group model with multifaceted benefits, but its success depends on the commitment of participants and their regular attendance.

### Scientific advisor participation

Coaches discussed SAs in 23 of the interviews. About half of these mentions came from coaches in the structured arm of the study, where SAs attended one group meeting, participated in a discussion session with just the participant and coach, and in some cases attended the mock review. From the perspective of those in the structured arm, benefits of the SAs included feedback from a subject matter expert and commitment to meeting attendance and participation. One coach mentioned that the combined feedback from the coach, peers, and SA helped to “validate” the ideas in the grant. Only one coach from the structured arm was critical of the SAs, noting that they did not participate in discussions or were defensive about the scientific merit of the participant’s project. Among those in the unstructured arm, coaches commented that SAs did not play a meaningful role because they were not involved directly in the coaching sessions. These coaches were also not clear on how much interaction the participants had with their SAs. In general, including SAs in the coaching group sessions can lead to a well-rounded coaching experience, where advisors’ subject matter expertise complements the coaches’ focus on grant writing skills.

### Mock review

Coaches discussed the mock review sessions in 31 interviews. In general, coaches found the mock review sessions to be a useful supplement to the group sessions and a way of rounding out the coaching experience with reviewers reading the proposals for the first time. Benefits included the simulation of a study section and feedback from a subject matter expert. Most coaches described the reviews as being highly rigorous, with occasionally “brutal” but constructive feedback that ultimately benefited the writer. Only rarely did coaches describe the sessions as unhelpful due to a lack of constructive or sufficiently critical feedback. More often, coaches noted that participants needed more guidance in selecting reviewers, and that participation was low due to participants’ lack of preparation or progress. Another concern was that, on occasion, participants chose reviewers from their own mentoring teams, and so the reviewers had a vested interest or bias in favor of the participants. Mock reviews can be extremely beneficial for grant writers, rounding out their experiences with critical, substantive feedback, so long as participants are sufficiently prepared and select experienced reviewers with subject matter expertise.

### Readiness to write

To establish eligibility for the study, the application included questions and documentation that the person was “ready to write”. However, it was not always easy to make that determination, and we tended to err on the side of inclusion. Coaches in 17 interviews commented on the problem of including participants who were not “ready” to write a grant, or those whose circumstances limited the feasibility of completing a grant proposal. In the words of the coaches, a lack of readiness meant that a participant did not have sufficient preparation, was not at an appropriate career stage, and/or lacked the institutional resources and mentorship to complete a competitive grant. Specific cases include lack of development of a grant idea, lack of institutional support for a mentored award, or lack of institutional resources to conduct the actual science. Coaches who worked with participants in these situations noted that this was a serious problem and affected how well the group worked in terms of the effectiveness and actionability of peers and coach feedback. Even though coaching groups are intended to systematize access to grant writing support, an assessment of writing readiness in terms of institutional resources and developed ideas is still important in selecting participants for this type of intervention.

### Mix of mechanisms

Twenty-six interviews mentioned the benefits and/or drawbacks of grouping participants working on a variety of grant mechanisms. In most cases, coaches described the mix of mechanisms as a challenge, noting difficulties with calibrating discussions around R-type and K-type awards, which contain different elements. Less frequently, coaches described the mix as a strength, noting that group members with experience with multiple grant mechanisms provided valuable feedback to their less experienced peers. These coaches also noted that the mix of mechanisms provided the opportunity for group members to learn more about the various grant types, knowledge which may benefit them in the future. In general, coaches prefer groups with participants working on similar mechanisms, or at least broadly grouped by R-type and K-type awards, but the potential benefits of mixed groups are notable.

### Mix of disciplines

Despite best efforts to create coaching groups comprising participants and coaches who worked in broadly similar research fields, coaches mentioned in 30 interviews that they and their participants worked across disparate disciplines. Coaches commented on the benefits and drawbacks of group misalignments. The strength was in preparing participants to write for reviewers with little to no knowledge of their specific research area. From one coach: “You need sets of objective, intelligent eyes looking at your document that aren’t part of your field.” The drawbacks were the lack of subject matter expertise and the inability to provide substantive feedback on the scientific aspects of the grants. Some coaches found it difficult to understand the science in the proposal and could only provide feedback on the structural components of the grants. By design, the coaching groups are not meant to provide extensive feedback on the science itself – participants receive that from their institutional mentors or from SAs and mock reviewers. Nevertheless, coaches were sometimes frustrated that they could not provide sufficient feedback on an individual participant’s science.

### Experiences of coaches across multiple groups

Three coaches who oversaw more than one group gave multiple interviews. Coaches described adapting their practices – or developing strategies to do so in the future – based on the situations of individual groups and their experiences across multiple groups. Coaches commented on how they adjusted (or would adjust) their practices based on the overall readiness of participants in the groups, for example by familiarizing participants with the NIH award process or adjusting the coaching schedule to focus additional time and energy on specific components of the grants where participants needed the most help, typically the Specific Aims page.

## Discussion

Perspectives provided by participants and coaches showed a strong convergence on several key findings about features that contributed to, or limited, the success of the trial’s five-month group coaching model employed to support grant proposal development. These findings include the high value of skilled coaching, the benefits of peer feedback and support, the importance of regular meetings with clear expectations and structured feedback, and the added value of SA participation and mock reviews. Participants and coaches alike acknowledged the overall value of the writing groups in supporting grant writing skill development, but they also identified areas that can limit the effectiveness of group coaching, particularly group composition and participants’ writing readiness.

### Critical role of coach experience and expertise leading the groups

A central theme emerging from participant interviews was the crucial role of coaching expertise in shaping successful experiences. Participants praised the coaches’ real-world knowledge of NIH-style grant review, their organizational skills, and their ability to provide both technical and motivational support. Coaches similarly viewed their role as pivotal but reported variation in coaching styles – ranging from strict grantsmanship to more holistic, socioemotional models. As noted by coaches who led more than one group, they adjusted their styles to the unique character of each group, pointing out the importance of training coaches in the group design. To improve consistency across coaches, study leaders joined early coaching groups of new coaches to provide feedback on their approach to the group coaching model. Those considering replicating this group model should select for faculty willing to put in this level of effort and they should be guided in the differences between the group model and feedback they are accustomed to giving on individual proposals.

The current mentoring literature emphasizes the importance of psychological safety in mentoring and peer interaction for providing open, confidential, and nonjudgemental spaces [13]. Peña and colleagues recently showed how even a short workshop based on social-emotional learning was able to shift participants from negative/challenge-focused attitudes and perceptions toward grant writing toward positive/process-focused ones [14]. Our findings support the idea that emotional safety and skill development need not be mutually exclusive and are beneficial when integrated effectively.

A recently reported study based on interviews with coaches who participated in one of the sites funded by NRMN observed very similar findings about the role of coaches and their perceptions of the important elements of coached writing groups [15]. Coaches emphasized the importance of creating a positive supportive environment, individualized feedback while writing, peer support, and normalizing the challenges writers were facing. The coaches also noted the importance of institutional support for success, and challenges managing groups that included individuals with a wide range of writing experience.

### Value of peer interactions and feedback

Peer interaction was another central theme that arose from the interviews. Participants valued the opportunity to give and receive feedback from colleagues at similar or more advanced career stages and across disciplines and institutions. These interactions often provided a sense of camaraderie and psychosocial support, and often improved writing clarity and increased confidence. The value of peer and near peer feedback is seldom appreciated and enabled in the traditional dyadic mentoring approach. This is another value added by the coached group model, in addition to the vicarious learning that takes place. The value of peer feedback was sometimes compromised in cases of poor attendance, uneven engagement, or mismatches in disciplinary expertise or grant mechanism within coaching groups. These limitations highlight the challenge of balancing diversity of perspective with alignment of experience.

### The role of scientific advisors with content/field expertise

One of the most novel contributions of this study is the evaluation of SA involvement, particularly in the structured arms of the study. Advisors who attended group sessions added valuable subject-matter expertise that complemented the mechanistic feedback provided by coaches and peers. In some instances, they strengthened mentor-mentee relationships. However, the variability in advisor engagement – especially in the unstructured arms – reveals the importance of formalizing expectations.

### Readiness to benefit from and contribute to a group

Coaches raised concerns about participants’ readiness to write and submit a grant, emphasizing that underdeveloped research ideas reduced the value of group participation. While participants did not consistently use the term “readiness,” some commented on their lack of preliminary data, the poor quality of the mentorship they receive at their home institutions, and the lack of institutional resources to support grant writing. Even well-designed coaching interventions cannot compensate for the absence of local support structures and research being insufficiently developed to form the basis of a proposal.

Inclusion of individuals at various stages of writing was purposeful in the study design to get a better idea of whom the model could benefit. Based on application materials, a determination of writing readiness was made that encompassed a range of experience levels, from being very advanced and likely to complete a proposal in five months, to those who had enough to start writing but for whom a submission could take up to a year (six months beyond completion of the initial five months of coaching). Since half were randomized into extended coaching for an additional 18 months, this longer time frame was well within the study parameters. But the design of requiring progress every two weeks toward a complete proposal turned out to be unrealistic for those just getting started. The group coaching model can be varied to accommodate different stages of writing, particularly with respect to the speed of expected progress. For example, multiple sessions can be allocated to the iterative refinement of the Specific Aims page, progressing to subsequent sections when it is final. This is a common approach of grant writing groups to specifically bring in those just getting started. But variations like this require a more relaxed pace and more extended group or individual coaching; a conscious decision can be made to include only those more advanced, less advanced, or a combination. Keeping expectations realistic for the stage of research and writing development is critical to ensure that those not as far along do not interpret taking more time as failure.

### Factors impacting writing progress during the group

In addition to understanding the attributes that affected coaching group efficacy, an essential goal of the qualitative analysis was to gain insights into why some individuals achieved proposal submission by the end of the five-month coaching groups while some did not. The data revealed a combination of potential explanatory factors, some related to individual participants, and some related to the efficacy of their group coaching experience. The data provide evidence that the variation across individuals was probably greater and more impactful than the variations across groups. For those who were seen by coaches as not ready to write, the limitations in their readiness were usually insurmountable during the five months of group work. Some of these individuals were likely able to continue working on their proposals, with or without the help of their coach, and achieve submission. Although both participants and coaches identified operational and group dynamic variations that could make groups efficacious, few instances were noted where difficulties within the groups rose to the level of being insurmountable. Additionally, as already noted, for those individuals just getting started on their proposals, the goal of finishing a new section every two weeks was not realistic, substantially lowering their chances of submitting at the end of the five-month period. For some, this was enough to cause them to disengage and stop coming to the group.

### Limitations

The interview data are robust and capture the commonalities of experiences among participants and coaches while also reflecting important variations in implementation of the coaching group model. The data provide some insights into the role of SAs in the model but they were not robust enough to reveal any systematic differences between structured and unstructured SA engagement. They also cannot provide insights into outcome differences between regular and extended dose as those would come from the analysis of interviews after the extended dose. Those data will be the subject of future publications analyses.

One potential limitation of the findings derives from the purposeful national study design. Some findings may not completely generalize to grant writing groups implemented at a single institution. Group dynamics could be different when peers share the same institutional context and coaches are known to the participants. However, from the personal experiences of the authors of this study, many of the complications of the national model are less challenging in an institutional setting. Rapport is usually easier to establish, peer relationships can enhance the process (very few instances of peer competition have been encountered), and coaches have more extended commitment to the individuals. With an institutional model, access to coaches after the group (like the extended dose) will be available as needed.

## Conclusions

Taken together, the results suggest that an optimal grant writing coaching group design should contain the following elements: (1) coaches who can draw on a high level of expertise from years of submitting and reviewing proposals; (2) coaches who actively support the structure of the grant writing groups and set clear expectations for participants; (3) coaches and participants who are reasonably well-aligned in their scientific fields and methodologies; (4) concordance among participants in their target grant mechanism (e.g., R versus K); (5) coach and participants working together to foster an environment with strong peer psychosocial support and feedback; (6) availability of a scientific advisor (e.g., mentor or field expert) to complement what is provided by the coaching group; and (7) alignment with participants’ writing readiness to benefit from the group design and expectations.

## Acknowledgements

This work was supported by funding from NIH, grant number U01GM132366 to Kola Okuyemi.

